# PLM_Sol: predicting protein solubility by benchmarking multiple protein language models with the updated *Escherichia coli* protein solubility dataset

**DOI:** 10.1101/2024.04.22.590218

**Authors:** Xuechun Zhang, Xiaoxuan Hu, Tongtong Zhang, Ling Yang, Chunhong Liu, Ning Xu, Haoyi Wang, Wen Sun

## Abstract

Protein solubility plays a crucial role in various biotechnological, industrial and biomedical applications. With the reduction in sequencing and gene synthesis costs, the adoption of high-throughput experimental screening coupled with tailored bioinformatic prediction has witnessed a rapidly growing trend for the development of novel functional enzymes of interest (EOI). High protein solubility rates are essential in this process and accurate prediction of solubility is a challenging task. As deep learning technology continues to evolve, attention-based protein language models (PLMs) can extract intrinsic information from protein sequences to a greater extent. Leveraging these models along with the increasing availability of protein solubility data inferred from structural database like the Protein Data Bank (PDB), holds great potential to enhance the prediction of protein solubility. In this study, we curated an Updated *Escherichia coli* (*E.coli*) protein Solubility DataSet (UESolDS) and employed a combination of multiple PLMs and classification layers to predict protein solubility. The resulting best-performing model, named Protein Language Model-based protein Solubility prediction model (PLM_Sol), demonstrated significant improvements over previous reported models, achieving a notable 5.7% increase in accuracy, 9% increase in F1_score, and 10.4% increase in MCC score on the independent test set. Moreover, additional evaluation utilizing our in-house synthesized protein resource as test data, encompassing diverse types of enzymes, also showcased the superior performance of PLM_Sol. Overall, PLM_Sol exhibited consistent and promising performance across both independent test set and experimental set, thereby making it well-suited for facilitating large-scale EOI studies. PLM_Sol is available as a standalone program and as an easy-to-use model at https://zenodo.org/doi/10.5281/zenodo.10675340.

## Introduction

Proper folding of proteins to maintain enough solubility and homeostasis is essential for nearly every protein-based biological process. Unsatisfied solubility or aggregation can impede protein-based drug development, such as antibody production. The low solubility of antibodies may limit their shelf-life and potentially induce adverse immune responses (1–3). Apart from antibodies, more and more enzymes of interest (EOI) are being discovered with an increasing speed, due to the decreasing cost of sequencing and gene synthesis as well as continuous improvement of high-throughput functional screening platforms (4–6). In these large-scale EOI screening studies, enhancing the accuracy of protein solubility prediction can improve the success rate of protein purification and facilitate the downstream biophysical or biochemical characterization. Common hosts such as bacterial cells, insect cells, yeast cells, plant and mammalian cells are often used for recombinant protein expression (7). Among these options, bacterial cells, typically *Escherichia coli* (*E. coli*), provide the advantage of easy genetic manipulation and cost-effectiveness, therefore serving as one of the major platforms for recombinant protein production (8). Improving the accuracy of protein solubility prediction in *E. coli* thus has great potential to reduce experimental cost and increase the success rate of novel EOI discovery.

Protein solubility in *E. coli* is a complex issue influenced by numerous factors at different levels. Firstly, regarding the sequence level, several attributes have been identified as pivotal determinants of solubility, encompassing the composition of specific amino acids (Asn, Thr, Tyr), the frequency of tripeptides (9) and the ratio of charged amino acids on the protein surface (10,11). Secondly, during the protein expression process, the ineffective translation of the mRNA and the manifestation of protein toxicity to *E. coli* may impede normal growth. Additionally, there is evidence suggesting that proteins instability can lead to aggregation, thereby inducing the formation of inclusion bodies (12–14). Thirdly, at the experimental level, factors such as the selection of expression strains, appropriate fusion tags, culture temperature, and the pH value of the protein lysis buffer collectively contribute to the protein solubility (15). Notwithstanding the abundance of available data, extracting essential features responsible for insolubility remains difficult, rendering the prediction of protein solubility a challenging task.

Over the past three decades, extensive researches have been dedicated to investigate the correlation between protein sequence and solubility, culminating in the formulation of numerous predictive models. Early studies employed statistical methods with a small dataset to extract essential protein sequence features for solubility classification (16). Additionally, SWI (17) employed the arithmetic mean of sequence composition scoring to predict protein’s solubility. Another booster for protein solubility predictor is the emergency of machine learning (ML), which shifted the focus towards the utilization of feature engineering and supervised ML algorithms, such as linear regression, support vector machines, and gradient boosting machines (18). Some of the models leveraging traditional ML algorithms, including PROSO (19), SOLpro (20), PROSO II (21), SCM (22), Protein-Sol (23), PaRSnIP (24) and SoluProt (25). Recently, with the fast development of deep learning (DL), a transition from conventional ML to DL algorithms has been observed (26). Researchers generated several DL models, such as DeepSol (27), SKADE (28), EPSOL (29), DSResSol (30) and DeepSoluE (31), which employ Convolutional Neural Networks (CNNs), bidirectional Gated Recurrent Unit (biGRU) or Long Short-Term Memory (LSTM)-based approaches for protein solubility prediction.

With the continuous development of natural language processing (NLP), attention-based algorithms deepen the understanding of relationships among tokens (32,33). The protein sequences, serving as the language of proteins, have catalyzed the development of diverse protein language models (PLMs), such as ProteinBERT (34), Evolutionary Scale Modeling (ESM) (35,36), and ProtTrans (37). These models excel in general contextual embeddings of protein sequences by training transformer models on large protein databases, such as UniRef (38) and BFD (Big Fantastic Database) (39). By training with a labeled dataset, these models can be fine-tuned for predicting specific protein properties. For example, NetSolP utilized ESM1b and employed multilayer perceptron (MLP) for protein solubility classification (40).

Despite the significant advancements achieved by the aforementioned models, there still exists room for improvement. First of all, high-quality training data are critical for the success of any model. However, the most widely utilized dataset to date, originating from PROSO II (21) in 2012, has become outdated and suffers from ambiguous annotations. Another frequently used dataset, provided by SoluProt (25) in 2021, comprises only 10,912 sequences. Secondly, the accessibility and usability of models are also critical. For example, some of the reported models are out of service (19,21,22), while others require the inclusion of protein secondary structure information as inputs which is time-consuming and compute-intensive (24,27,29,30). Thirdly, model architectures play a crucial role in DL-based classification tasks. With the expanding of Uniref50D (38) database, ESM2 updated multi-parameter versions of models (36), showcasing improvements in predictive capabilities and overall effectiveness in capturing intricate features of protein sequences. In terms of classification layers, the aforementioned NetSolP (40) applied only a simple MLP. Thus, the incorporation of updated protein language encoders and different classifier layers holds great potential to further improve prediction performance.

Given the aforementioned limitations, we attempted to improve the performance of protein solubility prediction by focusing on two key aspects: generating high-quality datasets, and developing a robust and easy-to-use model. For datasets generation, we integrated existing protein solubility data from TargetTrack (41), DNASU (42), eSOL (43) and Protein Data Bank (PDB) (44), making diligent efforts to improve the accuracy and comprehensiveness of the dataset. We designated this compiled dataset as the Updated *E. coli* protein Solubility DataSet (UESolDS). For model design, we trained a series of architectures by integrating multiple PLMs with diverse classifiers. Three pretrained PLMs, namely proteinBERT, ESM2 and ProtTrans were employed alongside different classification layers, comprising MLP, Light Attention (LA) (45) and the biLSTM_TextCNN (46). Subsequently, we systematically evaluated their performance on an independent test set. In comparison to previously reported models, the combination of ProtT5-XL-UniRef50 (ProtT5) with biLSTM_TextCNN demonstrated superior performance, denoting as the Protein Language Model-based protein Solubility prediction model (PLM_Sol). Finally, we performed an experimental test by assessing the solubility of 216 understudied proteins belonging to three distinct family types, and PLM_Sol consistently exhibited the best performance.

## Materials and methods

### UESolDS source

*TargetTrack* The TargetTrack database compiles experimental results curated by over 100 investigators across 35 centers from 2000 to 2015, focusing on investigating the expression levels and structures of 350,000 proteins (41). We applied a comparable data filtering analysis approach as that employed in SoluProt (25). Utilizing experimental protocols provided by each contributor in the TargetTrack database, we extracted 17 datasets specifically focusing on protein expression in *E. coli* (Supplementary Table S1). The TargetTrack database contained numerous entries marked as “work stopped” or “other” due to uncertain final statuses, and these data were consequently excluded (21). Subsequently, based on the experimental status of proteins, they were categorized into insoluble (Insol) and soluble (Sol). Insol proteins were those labeled with the tags “tested”, “selected”, “cloned” and “expression tested”. Sol proteins were those labeled with “expressed”, “soluble”, “purified”, “in PDB” and had subsequent structural analysis data.

*DNASU* The DNASU is a global plasmid repository. Data obtained from DNASU originating from the Protein Structure Initiative: Biology (PSI:Biology) (42). The entries using the common vectors (“pET21_NESG”, “pET15_NESG”) for recombinant protein expression were retrieved. Then they were categorized based on the following tags: Insol proteins were labeled as “Tested_Not_Soluble”, while Sol proteins were labeled as “Protein_Soluble”.

*eSOL* eSOL is a database offering solubility information for *E. coli* proteins through the protein synthesis using recombinant elements cell-free expression system (43). The eSOL database assigns a solubility score to Sol proteins, whereas those lacking solubility scores are categorized as Insol proteins.

*PDB* Protein sequences with *E. coli* as expression host from the PDB database (March 2023 version) were extracted and annotated as Sol (44).

### UESolDS data cleaning

Six steps were implemented sequentially for data cleaning to compile the aforementioned UESolDS.

1. Membrane proteins predicted by TMHMM (47) were filtered out, due to their commonly insolubility upon overexpression (21,25,48).
2. The following his tags fragments in proteins were excluded, due to an uneven distribution of these tags between Insol and Sol proteins revealed by NetSolP (40): “MGSDKIHHHHHH”, “MGSSHHHHHH”, “MHHHHHHS”, “MRGSHHHHHH”, “MAHHHHHH”, “MGHHHHHH”, “MGGSHHHHHHH”, “HHHHHHH”, “AHHHHHHH”.
3. Sequences containing special characters “X|x”, “U|u”, “Z|z”, “*”, “.”, “+”, “-” were thoroughly removed.
4. Sequences with lengths < 25 aa or > 2,500 aa were filtered out.
5. During the data categorization process, we identified data contamination, as some Insol proteins exhibit high similarity with Sol proteins. To enhance the dataset quality, we searched for matches of Insol protein sequences in Sol by using DIAMOND blastp, and removed overlap Insol entries with identity > 75% and coverage > 70%.
6. Clustering above filtered sequences using MMseqs (49) with 25% identity and coverage > 70% threshold.

Finally, there were 32,053 entries categorized as Insol and 47,291 entries categorized as Sol, resulting in the creation of the UESolDS.

### Independent test set generation

For an unbiased evaluation of model performance, we first aggregated the training sets of the previously reported models slated for evaluation in this study. Next, sequences in UESolDS with an identity of 25% or greater to the integrated training sets were excluded. Subsequently, we randomly selected 2,000 sequences from the remaining Sol/Insol data to form the independent test set.

### Model architecture and training process

*ProteinBERT* ProteinBERT has six encoder layers and is trained on the Uniref90 dataset, along with protein gene ontology annotation information. The input of proteinBERT are the protein sequences and the outputs are fixed-dimension vectors. Then the vectors are fed into a vanilla MLP layer to obtain solubility probabilities. For the training process, ProteinBERT underwent a process where all layers of the pretrained model were initially frozen, except for the classification layers, which were trained for up to 10 epochs. Afterward, all layers were unfrozen and trained additional epochs until the test loss did not decline for 3 epochs. To optimize the learning process, the dynamic learning rate adjustment technique ReduceLROnPlateau was employed. The loss was calculated using binary cross-entropy. The training process was executed on a single GPU (Tesla P100-PCIE-12GB, with the learning rate and batch size set to 0.001 and 80, respectively.

*ProtTrans* To employ ProtTrans as the encoder, six models were trained by combining two architectures from ProtTrans with three classifiers respectively. The first architecture, ProtBert_BFD, based on BERT with a total of 30 encoder layers, is trained on the BFD dataset. The second one is ProtT5, utilized an architecture of T5 with a total of 24 encoder layers. It is pretrained on the BFD dataset and fine-tuned on Uniref50. The inputs of these PLM models are the protein sequences, and the outputs are matrices whose shape is n×L, where n represents the embedding dimensions, and L is the maximum protein sequence length. Following the ProtBert_BFD and ProtT5 architectures, three classification modules were introduced and tested respectively. The first classification module is a vanilla MLP. The second is the LA architecture, as described in a previous study (45), which shows excellent performance in protein localization classification. Thirdly, the biLSTM_TextCNN architecture, known for its effectiveness in sentiment classification (46), utilizes a bidirectional LSTM (biLSTM) to convert the information from the encoder into corresponding matrices. Following the classification process, a Text-attentional CNN (TextCNN) is applied, featuring three parallel one-dimensional convolution (Conv-1D) layers for crucial feature extraction. Next, the max pooling is executed along the direction of Conv-1D. The resulting three vectors obtained are concatenated and feed into fully connected layers to generate probabilities. For the training process, Bio-embedding software (50) was used to extract embedding information from ProtBert_BFD and ProtT5. In the model training phase, all layers of the pre-trained model remained frozen. The classification layers were trained for 15 epochs. The learning rate was initially set to 0.001 and adjusted by the optimizer AdamW (51) with a weight decay of 0.001. Binary cross-entropy was utilized to calculate the loss. The training process was conducted on a single GPU (Tesla P100-PCIE-12GB) with a batch size of 72.

*ESM2* To employ ESM2 as the encoder, six models were trained by combining two architectures from ESM2 with three classifiers respectively. As for ESM2-based models, two implementations, esm2_t30_150M_UR50D (ESM2_30) and esm2_t33_650M_UR50D (ESM2_33), were selected. After the protein sequences are input into ESM2, the outputs matrices’ shape was L×n. L is the maximum protein sequence length and n is the embedding dimension. The classification modules utilized the same settings as the classification modules used above ProtTrans. For the training process, fine-tuning involved unfreezing the embedding norm before layers, embedding norm after layers, and Roberta Head for masked language modeling layers in the pretrained model. Training consisted of 15 epochs with a learning rate of 0.001 and a batch size of 119. The optimizer utilized AdamW (51) with a weight decay of 0.0001. Cross-entropy was used to calculate the loss. A total of 8 Tesla P100-PCIE-12GB GPUs were used for this process.

### Experimental test set generation procedure

To better understand PLM_Sol’s performance on the understudied EOI, we selected various types of in-house EOI, consisting of tandem repeat proteins, DNA transposases, and deaminases (unpublished data). These EOI were intended to be synthesized for the development of novel gene editing tools, without any prior solubility prediction steps. To minimize redundancy, we clustered these EOI at 25% sequence identity, yielding 216 entries including 155 tandem repeat proteins, 30 DNA transposases and 31 deaminases. Then we performed *in vitro* recombinant protein expression assay in *E. coli* using the following procedure.

The pET28a vector with an N-terminal His tag and SUMO tag was applied for protein expression in *E. coli* Rosetta (DE3) cells. Protein expression was induced by adding 0.1 mM IPTG when the OD600 reached 0.6-0.8, followed by incubation at 16 °C for 18 hours. Cell pellets were sonicated (50W, 3s on/3s off on ice for 3 minutes) after being resuspended in binding buffer (50 mM Tris-HCl, 500 mM NaCl, PH 7.5). Supernatant was extracted and analyzed via SDS-PAGE gels, where protein bands with overexpression strips at the right molecular weight were identified as Sol hits based on the Coomassie blue staining gel results.

### Evaluation metrics

The performance metrics, including accuracy, Area Under Curve (AUC), precision, sensitivity, specificity, F1_score, confusion matrix and Matthews Correlation Coefficient (MCC) were calculated using the scikit-learn packages (52).

## Results

### Dataset organization and analysis

The previously reported datasets are outdated, with ambiguous annotation (21) or limited coverage (25). To generate a more comprehensive and precise dataset, we reconstructed the dataset on recombinant protein solubility in *E. coli* by integrating and updating related databases, including TargetTrack, DNASU, eSOL and PDB. We executed a six-step filtration process, encompassing the removal of membrane proteins, His tag fragments, special characters, sequences within certain length ranges and ambiguous Insol sequences (Methods, Figure 1A). This refined dataset was denoted as the Updated *E. coli* Protein Solubility DataSet (UESolDS) with a total of 79,344 entries. In order to establish an independent test set, we conducted a sequence identity analysis between UESolDS and the integrated training datasets of seven previously reported models, including Protein_sol (23), SKADE (28), SWI (17), Soluprot (25), EPSOL (29), NetSolP (40) and DeepSoluE (31). Sequences with more than 25% identity to any of the previously reported training sets were removed. Subsequently, a random selection of 2,000 Sol and 2,000 Insol entries from the remaining dataset were compiled to form the independent test set (Figure 1A).

**Figure 1.**
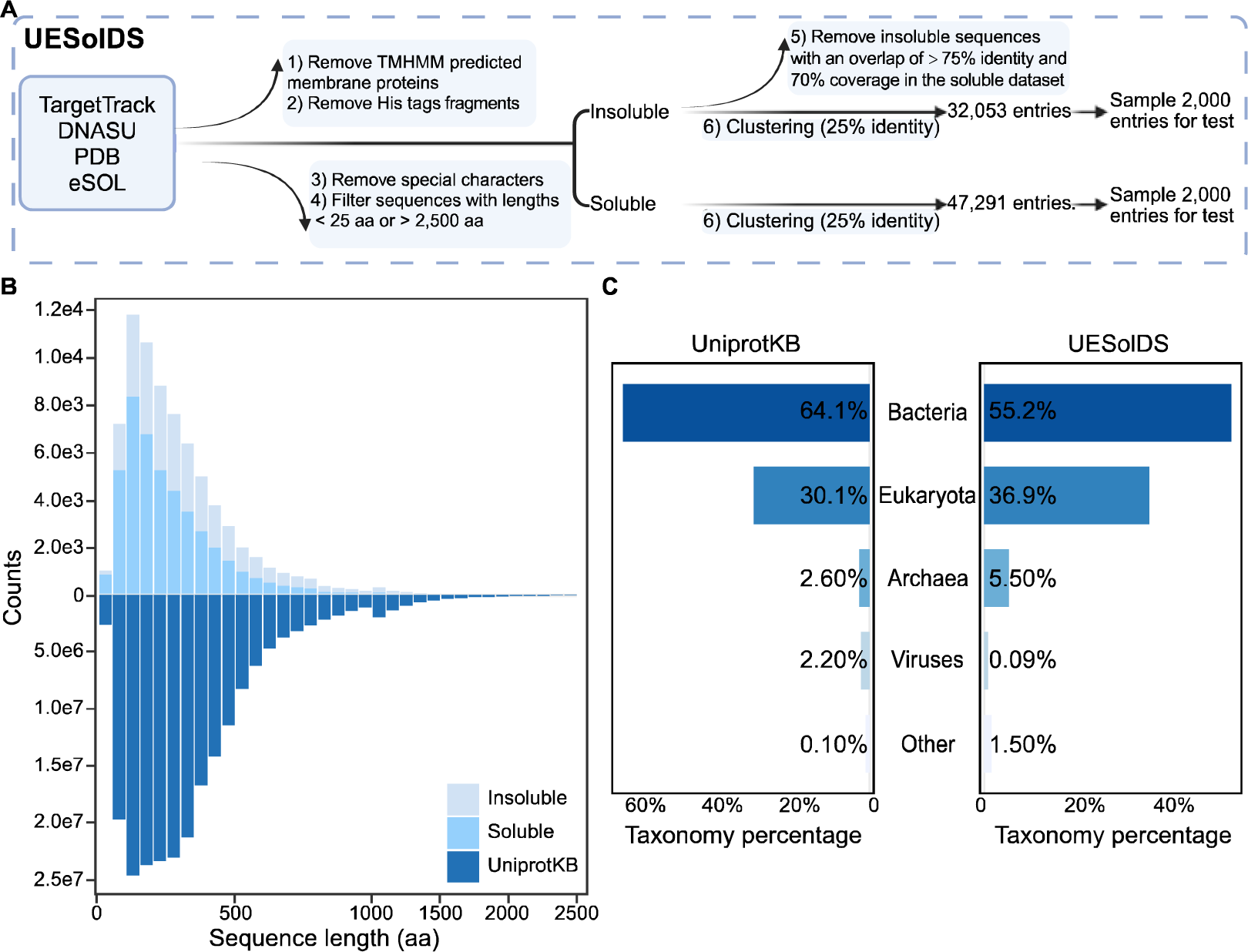
Generation and analysis of UESolDS. (A) Data processing pipeline of UESolDS. (B) Distribution of sequence lengths in the UniProtKB (release 2024_01) and UESolDS. (C) Taxonomy distribution of sequences from the UniProKB and UESolDS.

To elucidate the characteristics of UESolDS, we conducted an analysis of the sequence length and species distribution. The results revealed a concentration of protein lengths within the range of 25-500 amino acids in UESolDS (Figure 1B). It’s worth noting that UESolDS showed a very similar distribution pattern with UniProtKB (53) database (Figure 1B), indicating a good coverage and representation of the existing protein space. The statistical analysis of the species distribution showed that bacteria constituted the majority with about 55%, followed by eukaryotes with about 37%, archaea with about 5.5%, viruses with 2% and other sequences with 0.5%, which were annotated as unclassified (Figure 1C). Additionally, we compared UESolDS with UniProtKB, revealing a similar distribution in species composition, with bacteria, eukaryotes and archaea ranking as the top three categories (Figure 1C). These findings collectively emphasized a broad diversity of our refined dataset, thereby establishing a solid foundation for improving the model’s generalization capabilities.

### Model construction and performance evaluation

With the rapidly evolving development of PLMs, the utilization of updated PLMs like ESM2 (36), coupled with different classification layers holds the potential to improve the extraction of protein solubility features with higher accuracy. Therefore, we constructed models based on the following two parts: an encoder responsible for generating protein sequence embeddings from the PLMs, and a classification module for selecting and categorizing the crucial features. In this work, three PLMs have been tested, including ProteinBERT (34), ProtTrans (37) and ESM2 (Table 1). Additionally, three classification architectures have been employed, namely MLP, LA (45) and biLSTM_TextCNN (46). A total of 13 models were developed and trained (Methods).

**Table 1.**
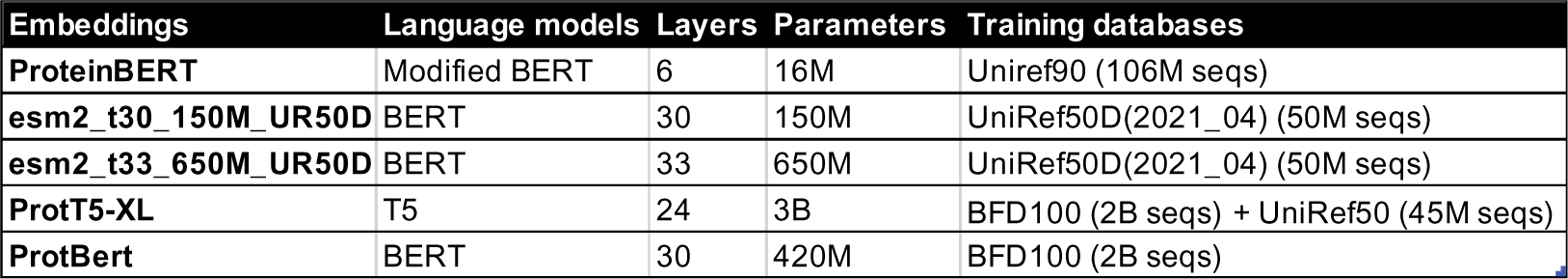
Characteristics of the PLM embeddings used in this study.

| Embeddings | Language models | Layers | Parameters | Training databases |
| --- | --- | --- | --- | --- |
| ProteinBERT | Modified BERT | 6 | 16M | Uniref90 (106M seqs) |
| esm2_t30_150M_UR50D | BERT | 30 | 150M | UniRef50D(2021_04) (50M seqs) |
| esm2_t33_650M_UR50D | BERT | 33 | 650M | UniRef50D(2021_04) (50M seqs) |
| ProtT5-XL | T5 | 24 | 3B | BFD100 (2B seqs) + UniRef50 (45M seqs) |
| ProtBert | BERT | 30 | 420M | BFD100 (2B seqs) |

We then evaluated the performance of the 13 in-house trained models. Among these, the combination of ProtT5 with biLSTM_TextCNN demonstrated the best performance, achieving the highest accuracy of 0.7236, top F1_score of 0.7541, and maximum MCC of 0.4616 (Figure 2A, B, C). Hence, the combination of ProtT5 with biLSTM_TextCNN was selected as our protein solubility classification model, designated as the Protein Language Model-based protein Solubility prediction model (PLM_Sol) (Figure 3A).

**Figure 2.**
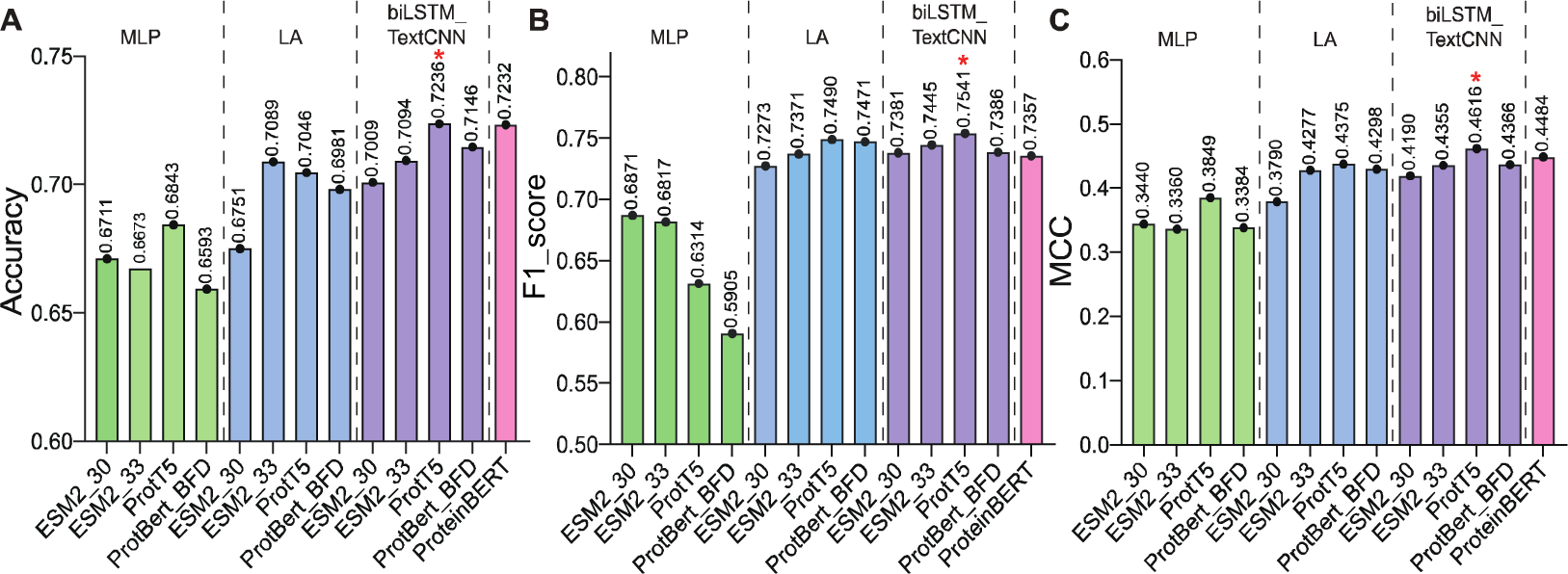
Evaluation of test set performance among various in-house models. (A) Comparison of accuracy among 13 in-house models. The green, blue and purple columns showed the results of different PLMs combined with MLP, LA and biLSTM_TextCNN classifier, respectively. The last pink column is ProteinBERT combined with MLP. (B) Comparison of F1_score among 13 in-house models. The assigned colors remain consistent with the annotation in (A). (C) Comparison of MCC among 13 in-house models. The assigned colors remain consistent with the previous annotation in (A).

**Figure 3.**
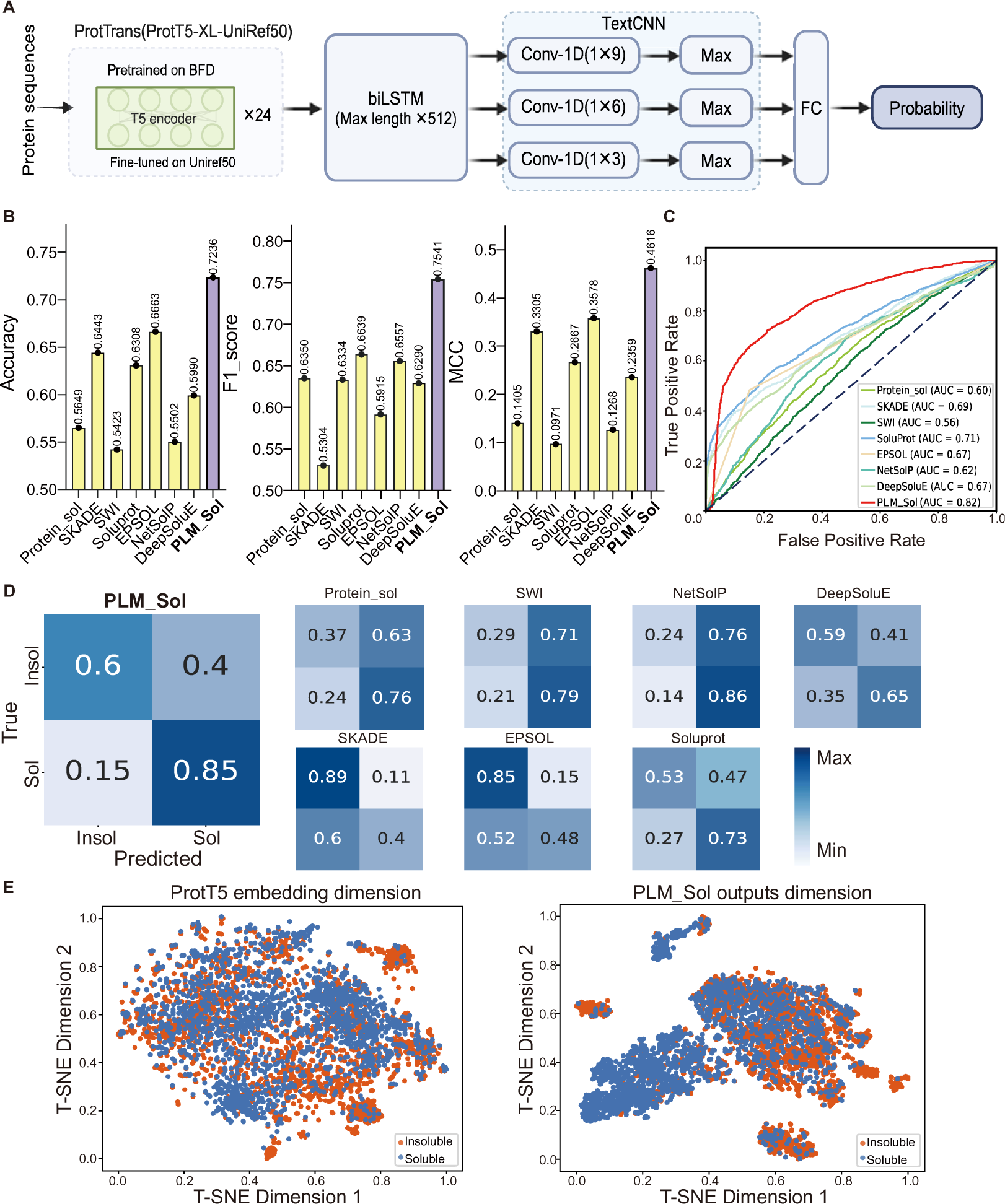
A) Model architecture of PLM_Sol; FC, fully connected layer. B) Comparison of accuracy, F1_score, and MCC between PLM_Sol and previously reported models. The yellow and purple columns represent the results of previously reported models and PLM_Sol, respectively. C) ROC curve for PLM_Sol and previously reported models. D) Confusion matrices depicting PLM_Sol predictions in comparison with previously reported models on independent test set. E) Dimensionality reduction using t-SNE for ProtT5 embeddings and PLM_Sol outputs on test set. Each point represents a sequence.

Subsequently, we compared PLM_Sol with seven previously reported models, and PLM_Sol exhibited enhancement over the previously best-performing software across various metrics. For example, PLM_Sol showed a 5.7% increase in accuracy over EPSOL, a 9% increase in F1_score over Soluprot and a 10.4% increase in MCC score over EPSOL on the test set. (Figure 3B, Table 2). PLM_Sol also showed the highest AUC score among the models evaluated based on the Receiver Operating Characteristic (ROC) curve (Figure 3C). Next, we analyzed the confusion matrix of each models and observed distinctive prediction outcomes among the previously reported tools (Figure 3D). For example, Protein_sol, SWI and NetSolP preferred to predict proteins as Sol, while SKADE and EPSOL preferred to predict proteins as Insol. Conversely, PLM_Sol showed more accurate predictions power.

**Table 2.** Comparison of solubility prediction performance of PLM_Sol with existing models on an independent test set. T represents the threshold value of the models. The bold string indicates the highest score within the respective column.

| Models | T | AUC | Accuracy | F1_score | MCC | Precision | Sensitivity | Specificity |
| --- | --- | --- | --- | --- | --- | --- | --- | --- |
| Protein_sol | 0.5 | 0.5985 | 0.5649 | 0.6350 | 0.1405 | 0.5470 | 0.7569 | 0.3729 |
| SKADE | 0.5 | 0.6882 | 0.6443 | 0.5304 | 0.3305 | **0.7811** | 0.4015 | **0.8874** |
| SWI | 0.5 | 0.5597 | 0.5423 | 0.6334 | 0.0971 | 0.5284 | 0.7905 | 0.2938 |
| Soluprot | 0.5 | 0.7126 | 0.6308 | 0.6639 | 0.2667 | 0.6095 | 0.7290 | 0.5325 |
| EPSOL | -- | 0.6664 | 0.6663 | 0.5915 | 0.3578 | 0.7630 | 0.4830 | 0.8499 |
| NetSolP | 0.5 | 0.6183 | 0.5502 | 0.6557 | 0.1268 | 0.5312 | **0.8564** | 0.2437 |
| DeepSoluE | 0.4 | 0.6660 | 0.6178 | 0.6290 | 0.2359 | 0.6110 | 0.6480 | 0.5875 |
| **PLM_Sol** | 0.5 | **0.8203** | **0.7236** | **0.7541** | **0.4616** | 0.6792 | **0.8474** | 0.5998 |

To visually showcase the classification performance of PLM_Sol, we applied t-SNE (54) to the test set using ProtT5 embeddings and PLM_Sol output data. Compared with the ProtT5 embeddings, the data processed by the PLM_Sol classification layer effectively separated Sol and Insol proteins data, suggesting a robust classification performance (Figure 3E).

### Solubility prediction for diverse types of EOI

To further validate the generalization of PLM_Sol, we conducted an evaluation of protein solubility prediction for 216 in-house EOI (Methods). These diverse protein types exhibit significant variability in solubility, some of them are reported to be poorly soluble (55,56), providing valuable material for the assessment of the model’s generalizability. We utilized the *in vitro* recombination protein expression assay to evaluate the solubility of EOI, resulting in 134 Insol proteins and 82 Sol proteins (Supplementary Table S2).

Next, PLM_Sol along with seven previously reported models were employed to predict the solubility of the aforementioned in-house EOI, followed by an assessment of their overall performance. The confusion matrix was utilized to showcase the comparison between predicted and experimentally observed outcomes (Figure 4A). Predictive results similar with previous models were observed. In contrast, PLM_Sol demonstrated a higher proportion of correct predictions, suggesting a better performance in predicting protein solubility for EOI screening (Figure 4B).

**Figure 4.**
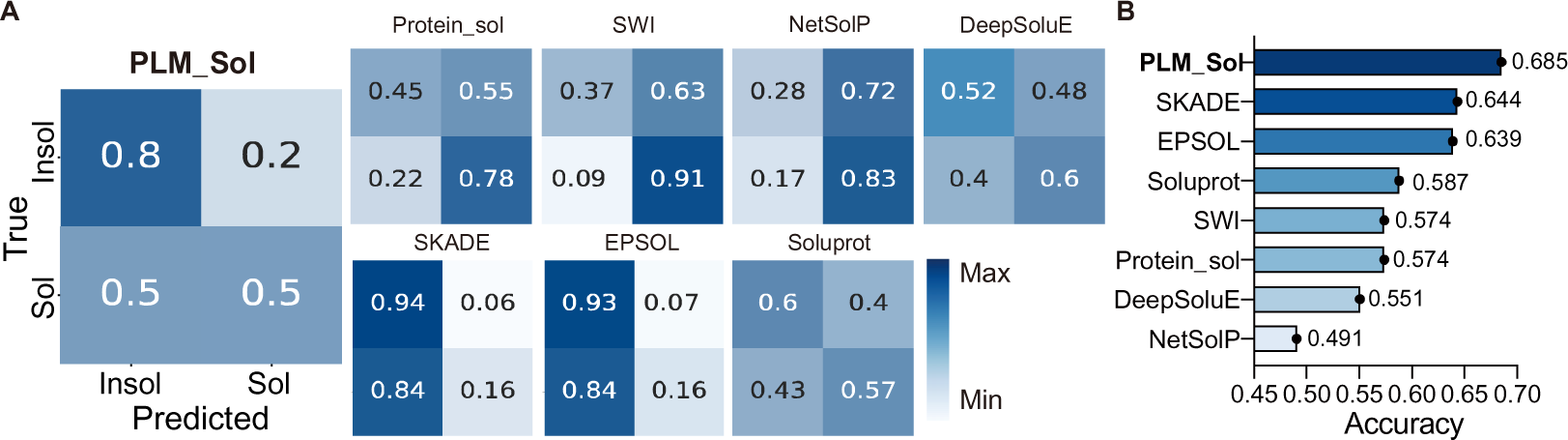
A) Confusion matrices depicting PLM_Sol predictions in comparison with previously reported models on the experimental test set. B) Accuracy for the tested models.

## Discussion

Deep learning relies on two fundamental pillars: data and model (57–59). However, in the development of solubility prediction tasks, most models have predominantly focused on model optimization, overlooking the quality of the datasets (27–29). For instance, a commonly used dataset from PROSO II (21) showed data contamination, as we identified 9.8% of Insol proteins that showed strong similarity (> 90% identity and > 70% coverage) to those present in the Sol dataset. This can be attributed to the ambiguity of the annotations for Insol proteins in the TargetTrack database, which serves as the dominant source of Insol data. In this study, a more comprehensive and accurate database UESolDS was generated by integrating the existing datasets, followed by a more stringent blastp parameters (>75% identity and > 70% coverage) to eliminate overlapping Insol instances within the Sol category. Furthermore, given the inherent complexity of protein insolubility states, more in-depth investigation and detailed annotation of insoluble protein experimental information may further improve the quality of the datasets. This, in turn, holds potential to further enhance the classification capability of DL models.

For model construction, we utilized the refined databases and employed PLMs incorporating additional classification layers for enhanced solubility prediction. After evaluation on an independent test set, we identified the best-performing model, named PLM_Sol. Compared to the state-of-the-art models, PLM_Sol exhibited a 5.7% increase in accuracy on the test set. In addition to the independent test set, we also validated the efficiency of PLM_Sol using our in-house experimental data, which showed improved prediction performance with an increase in accuracy of approximately 4.1%. Furthermore, our models could benefit from further optimization. Evaluation of the test set indicates there is room for improvement in specificity, which would subsequently enhance accuracy. In the fast-paced world of large language models, the utilization of parameter-efficient fine-tuning modules, such as LoRA (60), Adapter Tuning (61), IA3 (62) and Prompt Tuning (63), may enhance the model performance. Alternatively, employing the protein structure features derived from AlphaFold2 (39) or ESMFold (36), may provide the capability to identify key structural features to better separate Sol and Insol proteins. Improvement based on these continuously emerging algorithms, combined with updated high-quality datasets would ensure a better prediction accuracy.

All in all, PLM_Sol is easy-to-use and provides accurate predictions of protein solubility based solely on protein sequences as input. The integration of PLM_Sol into a high-throughput EOI screening pipeline offers the potential to avoid Insol gene synthesis, thereby improving the success rate and facilitating the scale-up screen process. Furthermore, the ongoing accumulation of experimental results from the above design, coupled with the continuously updated UESolDS holds the prospect of iteratively refining PLM_Sol, thereby continuously enhancing its overall predictive performance.

## Acknowledgements

We thank Duan Liu for his assistance in project management. We thank Computer Network Information Center, Chinese Academy of Sciences for kindly providing the computing clusters for training the deep learning model. We also thank Prof. Yong E. Zhang and Dr. Daqi Yu for their insightful comments and constructive suggestions.

## Author contributions

W.S. and H.Y.W. supervised the project. H.Y.W., X.C.Z., X.X.H. and W.S. conceived and designed the study. X.C.Z., T.T.Z, Y.L., C.H.L. and N.X. conducted the expression experiments. X.C.Z. and X.X.H. performed the computational analysis. H.Y.W., X.C.Z., X.X.H. and W.S. and wrote the manuscript.

## Data availability

The data underlying this article are available at https://zenodo.org/doi/10.5281/zenodo.10675340. Any additional information required to reanalyze the data reported in this paper is available from the lead contact upon request.

